# Growth of *Staphylococcus aureus* in the presence of oleic acid shifts the glycolipid fatty acid profile and increases resistance to antimicrobial peptides

**DOI:** 10.1101/2024.05.03.592415

**Authors:** Djuro Raskovic, Gloria Alvarado, Kelly M. Hines, Libin Xu, Craig Gatto, Brian J. Wilkinson, Antje Pokorny

## Abstract

*Staphylococcus aureus* readily adapts to various environments and quickly develops antibiotic resistance, which has led to an increase in multidrug-resistant infections. Hence, *S. aureus* presents a significant global health issue and its adaptations to the host environment are crucial for understanding pathogenesis and antibiotic susceptibility. When *S. aureus* is grown conventionally, its membrane lipids contain a mix of branched-chain and straight-chain saturated fatty acids. However, when unsaturated fatty acids are present in the growth medium, they become a major part of the total fatty acid composition. This study explores the biophysical effects of incorporating straight-chain unsaturated fatty acids into *S. aureus* membrane lipids. Membrane preparations from cultures supplemented with oleic acid showed more complex differential scanning calorimetry scans than those grown in tryptic soy broth alone. When grown in the presence of oleic acid, the cultures exhibited a transition significantly above the growth temperature, attributed to the presence of glycolipids with long-chain fatty acids causing acyl chain packing frustration within the bilayer. Functional aspects of the membrane were assessed by studying the kinetics of dye release from unilamellar vesicles induced by the antimicrobial peptide mastoparan X. Dye release was slower from liposomes prepared from cells grown in oleic acid-supplemented cultures, suggesting that changes in membrane lipid composition and biophysics protect the cell membrane against peptide-induced lysis. These findings underscore the intricate relationship between the growth environment, membrane lipid composition, and the physical properties of the bacterial membrane, which should be considered when developing new strategies against S. aureus infections.

## 1. Introduction

*Staphylococcus aureus* remains a global human health threat, and research into successful treatments of multidrug-resistant infections continues to be on the list of top priorities for the World Health Organization [1,2]. Thus, staphylococcal physiology, pathogenesis, and antibiotic development continue to be areas of active investigation. The characteristics of pathogenic bacteria are not adequately mimicked by growth in artificial media, and the idea that the host serves as growth medium with significant impact on staphylococcal pathogenesis has been revisited [3,4]. A striking example of the differences between *S. aureus* grown in vivo- and in vitro is the incorporation of straight-chain unsaturated fatty acids (SCUFAs) into membrane phospho- and glycolipids from a variety of sources including complex biological materials, free fatty acids, triglycerides, and cholesteryl esters [5,6]. Exogenous fatty acids are incorporated in *S. aureus* lipids via the FakAB system [6–8], which bypasses the fatty acid synthesis (FASII) pathway and compromises the effectiveness of the FASII-targeting antimicrobials [9,10].

The major glycerolipids of the *S. aureus* membrane are the phospholipids, phosphatidylglycerol (PG), lysyl-phosphatidylglycerol (LPG) and cardiolipin (CL), and the glycolipids monoglucosyldiglyceride (MGDG), and diglucosyldiglyceride (DGDG) [11]. A common SCUFA that has been shown to be incorporated into all the major *S. aureus* phospho- and glycolipids, from a variety of sources, is oleic acid (C18:1Δ9), the major SCUFA in the human body [12,13]. When *S. aureus* is grown in conventional media, the fatty acids are exclusively a mixture of straight- and branched-chain fatty acids saturated fatty acids (SCFA and BCFA, respectively), with PG32:0 containing an *sn*-1 anteiso C17:0 and *sn*-2 anteiso C15:0 being a major species [14,15]. However, at *S. aureus* infection sites, the incorporation of host SCUFAs and the concomitant production of membrane lipids containing these fatty acids has been demonstrated [6,15,16]. Lipidomic studies have shown that PG 33:1, which is esterified with C18:1 at the *sn*-1 position, and anteiso C15:0 at the *sn*-2 position, is a major lipid species when the organism is grown in media that contain SCUFAs in various forms [12].

Thus, the fatty acid composition of *S. aureus* growing at an infection site will differ radically from that grown *in vitro*. Given the critical role of fatty acids in membrane structure and function, we were prompted to undertake a biophysical investigation of the membranes and lipids of *S. aureus* grown in the presence and absence of SCUFAs.

The utilization of exogenous fatty acids from the growth environment in the synthesis of membrane lipids has potentially significant consequences for the physical state of the resulting membrane [17,18]. To a first approximation, the physical state of a biological membrane at a given temperature is a function of the lipid acyl chain composition. The lipid component of all biological membranes, including those of bacteria, appears to be regulated such that the overall membrane melting transition is complete just before the growth temperature is reached [19,20]. Thus, a well-controlled feedback cycle must exist between the fatty acyl composition and ambient temperature in all organisms that lack temperature control. In fact, it has been proposed that the physical state of the bacterial membrane functions as an allosteric regulator of membrane protein function [21].

*S. aureus* lacks the ability to synthesize SCUFAs and, in the absence of an exogenous fatty acid source, relies on terminally branched fatty acids to produce membrane lipids with the appropriate melting temperature, Tm [22]. The incorporation of SCUFAs into *S. aureus* membrane lipids poses a unique problem for the organism. Aqueous suspensions of phospholipids containing oleic acid undergo a melting transition close to or even below the freezing point of water [23], much below the optimal growth temperature for most bacteria. How then does *S. aureus* compensate for the presence of SCUFAs in the cytoplasmic membrane to allow optimal growth? And what are the physical consequences of a cytoplasmic membrane enriched in SCUFA lipids? The answers to these questions are of more than academic interest since the bacterial membrane bacteria is a principal barrier between the cell interior and the exterior medium. Its composition and material properties not only regulate membrane protein function and control entry of antibiotics, it is also the first site of attack for antimicrobial peptides and other components of the innate immune system [24].

Changes in the lipid composition of bacterial membranes often correlate with changes in the degree of polarization or anisotropy of the fluorescence emitted by the lipophilic dye 1,6-diphenyl-1,3,5-hexatriene (DPH) [25]. In pure fluid-phase lipid vesicles, a low degree of anisotropy generally reflects the high rotational freedom of the fluorophore surrounded by lipids with highly disordered acyl chains. In the gel phase, lipid acyl chain order is high, and, consequently, the measured fluorescence anisotropy of DPH is much higher in gel-than in fluid-phase liposomes [25,26]. In phase-separated liposomes with coexisting fluid and gel phases, DPH partitions approximately equally between the two phases [27]. For this reason, DPH is well suited to track lipid phase transitions in pure lipid vesicles and its use to characterize the phase behavior of membrane lipids and their mixtures is well established [25–28]. Based on these studies, low anisotropy values associated with DPH fluorescence are commonly interpreted as reflecting a high fluidity of the membrane in which the fluorophore is embedded, and DPH anisotropy measured on cells grown under a variety of conditions is often thought to correlate directly with membrane fluidity [29,30]. However, while the interpretation of DPH fluorescence anisotropy in pure lipid vesicles is straightforward, the behavior of DPH in complex biological membranes that contain a full complement of proteins and non-polar lipids is more complicated. Even in pure lipid vesicles the distribution of DPH within the bilayer is not uniform [28,31], and some substances will shift the equilibrium location of DPH within the membrane [31–34]. This altered fluorophore distribution can lead to changes in the measured fluorescence anisotropy that is entirely unrelated to the rotational freedom of membrane lipid acyl chains. Moreover, DPH can bind to specific membrane components or membrane-bound proteins, in which case the measured fluorescence anisotropy ceases to report on the state of the bulk membrane but rather reflects the micro-environment of the membrane in which the fluorophore is located [35].

In the present work, we addressed the compositional remodeling of the cytoplasmic membrane in response to exogenous lipids by using two complementary approaches. Rather than relying on a membrane-embedded fluorophore for reasons indicated above, we used differential scanning calorimetry (DSC) to investigate the mixing of lipid components in membrane fragments and whole lipid extracts from *S. aureus* grown with and without supplementation with oleic acid. We also used the well-studied antimicrobial peptide mastoparan X (MasX) [36,37] to assess the susceptibility of liposomes prepared from whole lipid extracts to membrane-active peptides as a function of supplementation cultures with exogenous lipids.

We found that the incorporation of oleic acid into the bacterial cytoplasmic membrane of *S. aureus* led to a concomitant increase in long-chain saturated and unsaturated glycolipid species that promote the formation of inverted lipid phases at temperatures well above the growth temperature. We also found that this response to oleic acid incorporation led to a much increased resistance to permeabilization by the antimicrobial peptide MasX at physiological temperature.

## 2. Material & Methods

### 2.1 Bacterial strain and growth conditions

The *S. aureus* strain JE2, derived from the community-acquired methicillin-resistant strain USA300, was used in the current study [12]. Bacterial cultures were grown in Tryptic Soy Broth (TSB) (BD, Difco, Franklin Lakes, NJ) in a shaking incubator (200 rpm) at 37 °C. Cultures of 500 mL were grown in 2.0 L Erlenmeyer flasks to mid-exponential phase using a 1% overnight starting culture. Where indicated, TSB was supplemented with 70 μM oleic acid (Sigma-Aldrich, St. Louis, MO) dissolved in ethanol. Cultures were harvested by centrifugation at 9,800 x g at 4 °C and washed twice with 0.9% NaCl.

### 2.2 Isolation of cytoplasmic membrane fragments

Isolation of cytoplasmic membranes was carried out as described in Wilkinson et al. (1978)[38]. After washing, 500 mL cultures were resuspended in 100 mL of hypertonic sucrose solution containing lysostaphin (Sigma-Aldrich, St. Louis, MO) for cell wall digestion (0.5 M Tris-HCl pH 7.5, 0.145 M NaCl, 0.05 M MgCl_2_, 1.0 M sucrose, 12.5 μg/mL lysostaphin). Cells were incubated at 37 °C for two hours followed by centrifugation and resuspension in 100 mL of 0.05 M Tris-HCl pH 7.5 containing 5 μg/mL DNase I with stirring for 30 minutes. Membranes were pelleted at 38,000 x g for 30 minutes at 4 °C and washed in ddH_2_O.

### 2.3 Lipid extraction and fractionation

Whole lipids were isolated using the Bligh and Dyer method [39]. After washing cells, 500 mL cultures were resuspended in 30 mL 0.9% NaCl. The suspension was placed in a separation funnel followed by addition of 37.5 mL chloroform and 75 mL methanol, and the funnel was shaken vigorously. The suspension was allowed to stand at room temperature for two hours prior to the addition of 37.5 mL chloroform and 37.5 mL ddH_2_O. The solution was again shaken vigorously, and layers were allowed to separate overnight. The bottom layer was filtered through anhydrous sodium sulfate in a fluted filter paper into a round bottom flask for rotary evaporation to dryness. Lipids were resuspended in 2 mL chloroform, transferred to a Teflon-capped glass tube, dried down under nitrogen, and stored at -20 °C until use. Lipid fractionation was carried out on a 1 cm diameter 1.5 g unisil silicic acid 100 - 200 column, as published previously [38,40]. The column was washed with redistilled n-heptane, diethylether and chloroform. Total lipids were resuspended in chloroform and added to the column. Neutral lipids were eluted with chloroform. The glycolipid fraction was eluted using first a 1:1 (v:v) chloroform:acetone solution followed by pure acetone. The phospholipid fraction was isolated using first a 1:1 (v:v) chloroform:methanol solution followed by pure methanol. Fractions were rotary evaporated, transferred to a Teflon-capped glass tube, dried down under nitrogen, and stored at -20 °C until use.

### 2.4 Chemicals

Synthetic lipids were purchased from Avanti Polar Lipids (Alabaster, AL). Carboxyfluorescein (CF) was purchased from Thermo Fisher (Thermo Fisher Scientific, Waltham, MA). Organic solvents (HPLC/ACS grade) were purchased from VWR (VWR International, Radnor, PA). Mastoparan X was purchased from American Peptide Co. (Sunnyvale, CA). A peptide stock solution was prepared by dissolving the lyophilized peptide in deionized water/ethyl alcohol, 1:1 (v/v) at a final concentration of about 200 μM, which was stored at -80 °C. The peptide concentration of the stock solution was determined by measuring the absorbance at 280 nm and using the molar extinction coefficient of tryptophan of 5600 M^−1^cm^−1^.

### 2.5 Preparation of Lipid Vesicles

Multilamellar vesicles (MLVs) were prepared by two different methods. 1. Lipids were dissolved in a 1:4 mixture of methanol and chloroform in a round-bottom flask. The solvent was then rapidly evaporated using a rotary evaporator (Büchi R-3000, Flawil, Switzerland) at 60^°^C. The lipid film was placed under vacuum for a minimum of 4 hours and hydrated by the addition of buffer at 80 ^°^C. The buffer composition was 20 mM MOPS, pH 7.5 containing 0.02% NaN_3_, passed through a Chelex 100 column, 50-100 mesh (Sigma-Aldrich, St. Louis, MO) to remove residual Ca^2+^ ions. Lipid concentrations were determined by the Bartlett phosphate method [41], modified as previously described [42]. 2. Alternatively, the lipid mixtures were dissolved in cyclohexane containing 1 % (v/v) methanol and frozen using a dry ice-acetone slurry. The solvent was sublimed under reduced pressure, the resulting sample dried under vacuum, and hydrated in Ca^2+^-free MOPS buffer. Both methods yielded equivalent results.

For CF efflux experiments, the lipid film was hydrated after thorough drying under vacuum using a buffer containing 50 mM CF (20 mM MOPS pH 7.5, 0.1 mM EGTA, and 0.02% NaN_3_, 50 mM CF). The suspension of MLVs was extruded 10 times through two stacked Nucleopore polycarbonate filters (Whatman, Florham, NJ) of 0.1 μm pore size using a water-jacketed high-pressure extruder from Lipex Biomembranes (Vancouver, Canada) at 70 ^°^C. After extrusion, CF-containing LUVs were passed through a Sephadex-G25 column (Sigma-Aldrich, St. Louis, MO) to separate the dye in the external medium from the lipid vesicles. The buffer used was 20 mM MOPS pH 7.5, containing 100 mM KCl, 0.1 mM EGTA, and 0.02% NaN_3_, which is iso-osmolar to the CF-containing buffer. The suspension was diluted in buffer to the desired lipid concentration and used for fluorescence measurements.

### 2.6 Kinetics of Dye Efflux

The kinetics of CF efflux, measured by the relief of self-quenching of CF fluorescence, were recorded in an Applied Photophysics SX.18MV stopped-flow fluorimeter (Leatherhead, Surrey, UK). The excitation was at 470 nm and the emission was recorded through a long-pass filter OG-530 (Edmund Industrial Optics, Barrington, NJ). The peptide concentration was 5 μM in all experiments.

The curves of carboxyfluorescein release as a function of time were characterized by a mean relaxation time (*τ*), as described before [43]. Briefly, the mean relaxation time is obtained from the integral [44,45],

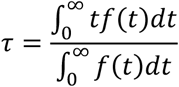

where

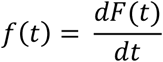

and *F(t)* is the experimental curve of normalized fluorescence increase as a function of time. This curve increases as CF is released, until it essentially reaches a plateau. The time-derivative of *F(t), f(t)* behaves as the probability density function [44,45]. For example, for a multi-exponential decay, *τ* is the weighted average of the relaxation times of each exponential function. Before numerical differentiation, the curves were smoothed as described before [43], to avoid errors due to experimental noise.

### 2.7 DSC

The thermotropic behavior of aqueous suspensions of membrane fragments and lipid extracts were analyzed using TA Nano DSC (Waters TA Instruments, New Castle, DE) and Malvern Panalytical MicroCal PEAQ-DSC (Malvern Panalytical Ltd., Malvern, UK) instruments. Three scans were performed on each sample under isobaric conditions using a scan rate of 30 °C/hour and temperature range of -3 °C to 80 °C. To prevent freezing, the suspensions and reference solution contained 30 % (v/v) ethylene glycol. To date, only heating scans on suspensions composed of MLVs have been analyzed. A baseline was obtained by fitting a polynomial function to the heating scans and subtracted to obtain the excess heating capacity. For ease of comparison, all DSC scans were normalized.

## 3. Results

### 3.1 DSC measurements

DSC has been used extensively to characterize the behavior of aqueous suspensions of pure lipid systems, their mixtures, but also that of more complex lipid preparations of biological origin [19,20,46,47]. DSC measures the heat capacity of the system under investigation by monitoring the amount of heat absorbed as the temperature is slowly raised, relative to a baseline. The result is plotted as the excess heat capacity, with peaks reflecting transitions between two states. Depending on the system studied, these can be associated with lipids, for instance, a lipid phase transition from gel to fluid, or a protein transitioning from a folded to a denatured state. The area under a peak is proportional to the enthalpy associated with that transition, and the width of the peak reflects its cooperativity. Complex mixtures of lipids and other membrane components like those investigated here can be expected to show generally broad transitions because the cooperativity of a phase transition is much reduced in multi-component mixtures, and the individual components will not mix ideally. Moreover, phase transitions in complex mixtures often show features, such as broad transitions broken up by peaks, which can indicate de-mixing of components, a phenomenon that is exacerbated by repeated heating.

Because of the complexity of biological membranes, we chose, wherever possible, to study a series of different preparations. The preparations that most closely capture the configuration of lipids and proteins in the intact cytoplasmic membrane are membrane fragments. We then subjected the isolated whole lipid components to DSC analysis, followed by the major lipid classes found in the *S. aureus* membrane, i.e., phospholipids, and glycolipids.

#### 3.1.1 Without fatty acid supplementation, *S. aureus* cytoplasmic membranes undergo a phase transition below the growth temperature

##### 3.1.1.1 Membrane Fragments

Membrane fragments contain polar and non-polar membrane lipids next to a full complement of membrane proteins. The DSC scans using membrane fragments revealed a phase transition that begins at 15 ^°^C and is complete at the growth temperature of 37 ^°^C (Fig. 1A). A smaller transition centered at 60 ^°^C was found to decrease with subsequent heating scans and is attributed to the irreversible denaturation of proteins in the sample (Fig. 1B), consistent with previous reports on whole *S. aureus* membrane preparations [19].

**Figure 1.**
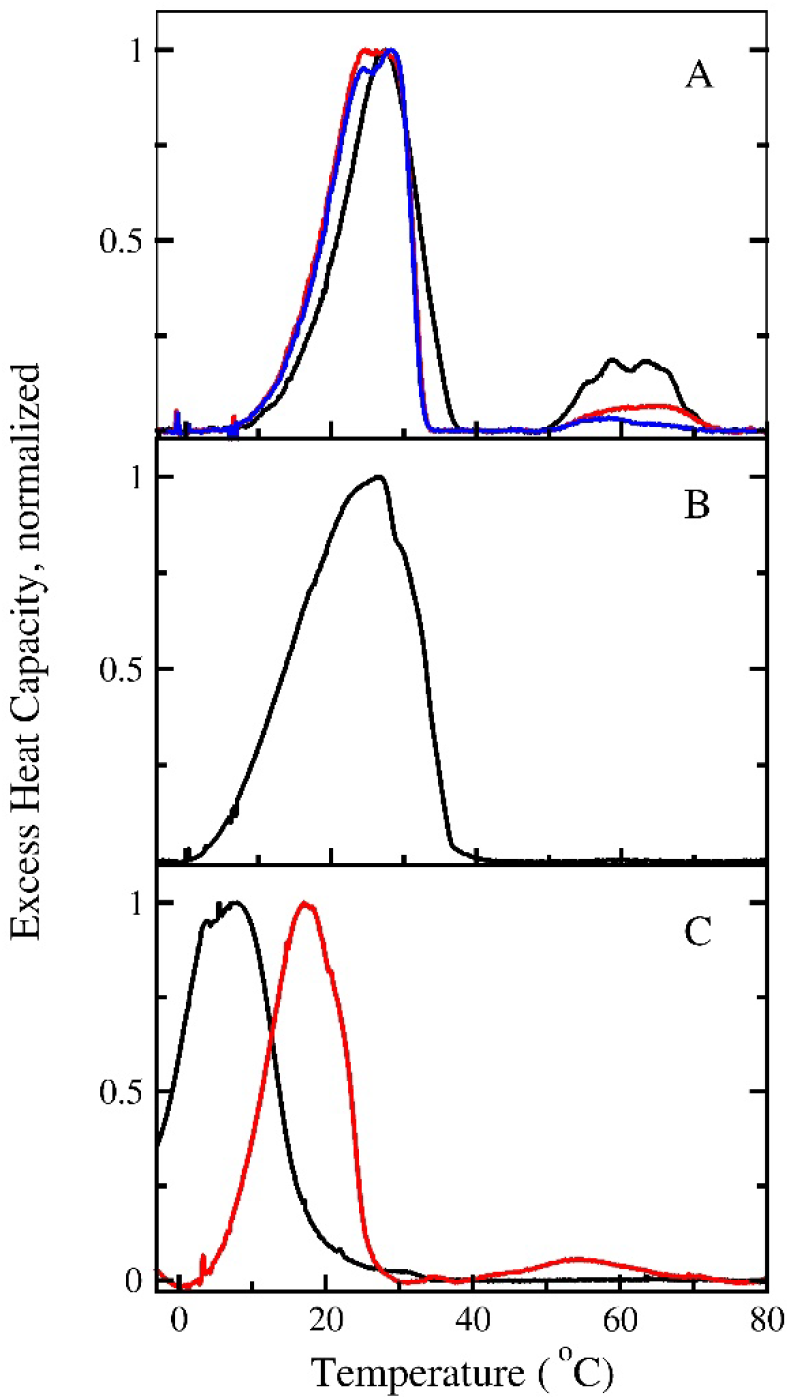
DSC scans of *S. aureus* membrane and lipid preparations from cultures grown in TSB without supplementation. Heating scans obtained from an aqueous suspension of membrane fragments (A). First heating scan, *black trace*; Second heating scan, *red trace*; third heating scan, *blue trace*. First heating scan of an aqueous suspension of MLVs made from whole lipid extracts (B). First heating scans of an aqueous suspension of MLVs (liposomes) made from the phospholipid and glycolipid fractions (C). Phospholipid fraction, *black trace*. Glycolipid fraction, *red trace*.

##### 3.1.1.2 Total Lipid Extracts

Total lipid extracts from cultures grown without supplementation show a single-phase transition below 40 ^°^C that is broadened with respect to the DSC scan from membrane fragments (Fig. 1 B). The observed broadening of this transition is likely due to a de-mixing of the lipid components in the sample during the preparation of the liposomes used for DSC, or the absence of protein in the lipid extracts. The high temperature transition observed in DSC scans of the membrane fragment sample (Fig. 1A) is missing in the total lipid extract preparation, which correlates with the absence of protein in the extracts.

##### 3.1.1.3 Phospholipid and Glycolipid fractions

In cultures grown in TSB without supplementation, PG and the glycolipids MGDG and DGDG are the most abundant lipid classes in the *S. aureus* cytoplasmic membrane [22,48]. In general, glycolipids show a chain melting transition from gel to fluid or to an inverted hexagonal phase that is shifted by 10-20^°^C to higher temperatures relative to the main transition in phosphatidylcholines or PGs with the same acyl chain composition [23]. The shift to higher melting temperature is due to the presence of hydrogen bonds between the headgroup sugar moieties of the glycolipids. We observed a similar difference in the main transition temperatures of the glycolipid and phospholipid fractions (Fig. 1C), which is supported by our previous observation that the fatty acyl composition is very similar in glyco- and phospholipids in cultures grown without supplementation [48].

#### 3.1.2 Membranes from cultures grown with oleic acid supplementation show a pronounced high temperature transition

##### 3.1.2.1 Membrane Fragments

The supplementation of culture medium with oleic acid leads to the appearance of membrane lipids with an odd number of carbons and unsaturated fatty acid chains, which is a clear indication that oleic acid is taken up from the medium and incorporated into membrane lipids [48]. In the DSC scans of membrane fragments that originate from oleic acid-supplemented cultures (Fig. 3 A), the minor transition that occurs below the growth temperature is shifted to lower temperatures with respect to the equivalent transition in unsupplemented cultures (Fig. 2). In oleic acid-supplemented cultures, the high-temperature transition is pronounced but, remarkably, does not disappear completely in repeated heating scans. In order to gain more insight into the nature of the high-temperature transition, we again analyzed total lipid extracts and phospholipid and glycolipid fractions in separate DSC scans.

**Figure 3.**
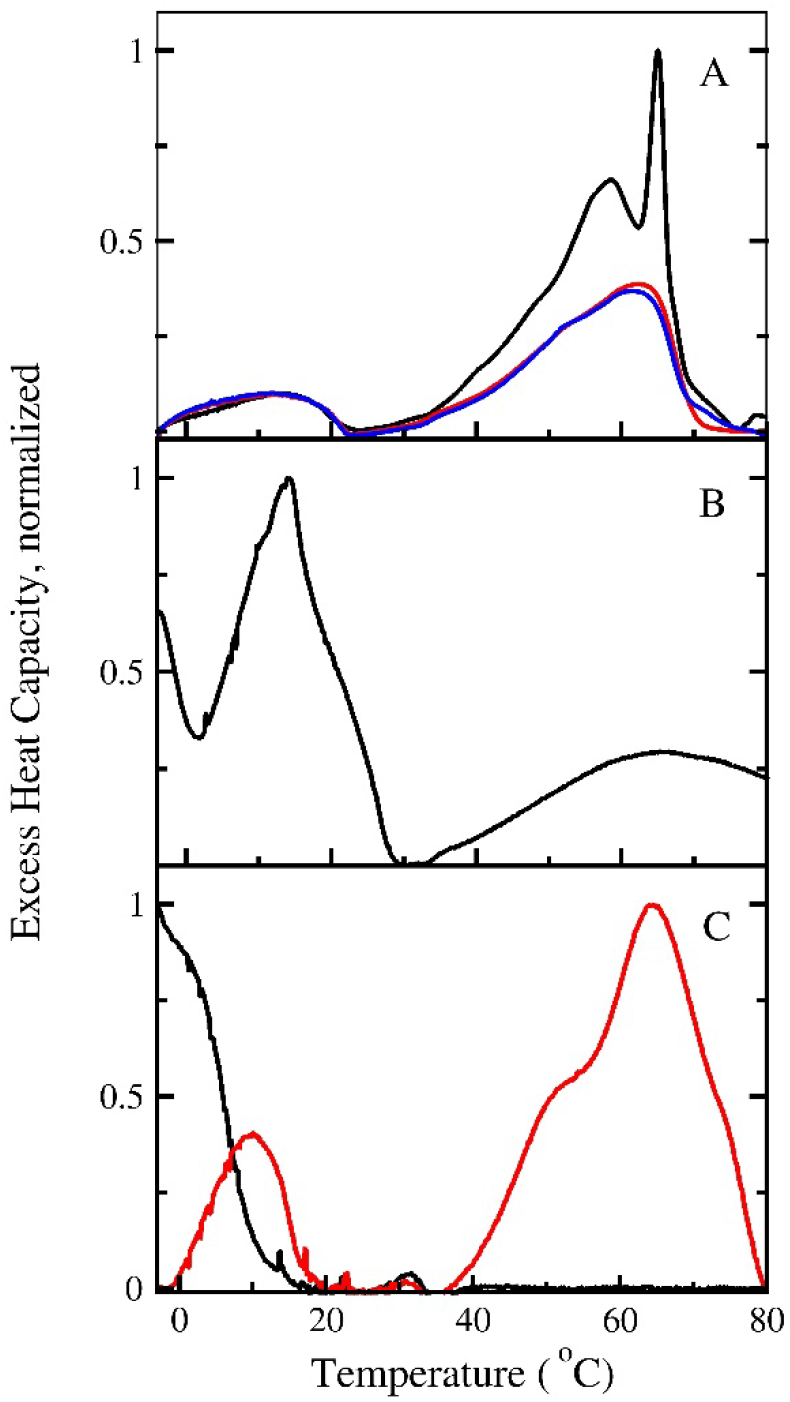
DSC scans of *S. aureus* membrane preparations from cultures grown in TSB with oleic acid supplementation. Heating scans obtained from an aqueous suspension of membrane fragments (A). First heating scan, *black trace*; Second heating scan, *red trace*; third heating scan, *blue trace*. First heating scan of an aqueous suspension of MLVs made from total lipid extracts (B). First heating scans of an aqueous suspension of MLVs made from the phospholipid and glycolipid fractions (C). Phospholipid fraction, *black trace*. Glycolipid fraction, *red trace*.

##### 3.1.2.2 Total lipid extracts

The DSC scans on total lipid extracts from oleic acid-supplemented cultures show a large transition below the growth temperature with a maximum between 10 and 20 ^°^C (Fig. 3 B). There appears to be an additional transition below 0 ^°^C, which suggests the presence of membrane lipids with unsaturated acyl chains that typically have gel-liquid crystalline transition temperatures below 0 ^°^C [23]. Moreover, the oleic acid-supplemented samples show a broad and featureless transition above the growth temperature, supporting the idea that the high temperature transition seen in the corresponding membrane fragment samples is not due to the presence of protein alone.

##### 3.1.2.3 Phospholipid and Glycolipid fractions

The most abundant PG species found in the cytoplasmic membranes of oleic acid-supplemented cultures have acyl chain compositions of 33:1, 33:0 and 35:1 and 35:0 [48]. The unsaturated species are most likely esterified with oleic acid in the *sn*-1 position, which is expected to shift the melting temperature of the PG fraction to a lower temperature compared to that isolated from cultures without the addition of exogenous fatty acids. Indeed, the phospholipid fraction isolated from oleic acid-supplemented cultures shows a phase transition that begins well below 0 ^°^C (Fig. 3C, compare Fig. 1C). In the glycolipid fraction from the same cultures, we found two transitions. The first of these is centered at 10 ^°^C and can be assigned to a glycolipid fraction of similar acyl chain composition as the phospholipid fraction that gives rise to the transition below 0 ^°^C. We again attribute the shift to a higher melting temperature in the glycolipid fraction compared to the phospholipid fraction to additional hydrogen bonding among the headgroup sugar residues in glycolipids. However, we also found an additional and pronounced high temperature transition between 40 and 80 ^°^C in the glycolipid fraction. This transition appears characteristic for lipids isolated from oleic acid-supplemented cultures and is absent in the corresponding fractions obtained from non-supplemented cultures. The appearance of the high-temperature transition in the glycolipid fraction suggests that the transition observed in whole lipid extracts and fragments from oleic acid-supplemented cultures is due to the presence of a glycolipid fraction with a fatty acyl composition that differs markedly from that melting around 10 ^°^C.

### 3.2 Kinetics of peptide-induced dye release from dye-encapsulated liposomes

Membrane-active peptides act at the bilayer-water interface of cytoplasmic membranes and liposomes without the involvement of surface receptors. In dye-encapsulated liposomes, binding usually leads to release of content at a rate that depends on the peptide and membrane lipid composition [49]. The majority of α-helical membrane-active peptides are between 15 and 35 amino acid residues long and carry a net-positive charge at physiological pH, which allows them to bind with high affinity to liposomes containing anionic lipids [49]. In many cases, the interaction of a membrane-active peptide occurs as a two-step process. The peptide first associates with the membrane surface, which we refer to as binding. The degree of surface binding is largely determined by the headgroup composition of the constituent lipids. In many cases, binding is followed by a second process during which the peptides become further embedded in the bilayer or cross it. This second step can lead to membrane disruption, sometimes transient, and is usually accompanied by release of content from liposomes [49]. The kinetics, or efficiency, of content release depends on the number of peptides bound per liposome, and the properties of the lipid bilayer, which is a function of the lipid acyl chain composition [49]. Peptide-induced release of content from liposomes is thus very sensitive to changes in lipid acyl chain composition [50] and an excellent means to assay membrane susceptibility to perturbation by external agents.

Here, we used Mastoparan X (MasX) to assess if the altered fatty acid composition in response to oleic acid supplementation has functional consequences on the susceptibility of lipid vesicles to a membrane-active peptide. The mastoparans, a class of mast cell degranulating peptides, were originally identified in the venom of *Vespa spp* [37,51]. MasX, isolated from the Japanese hornet *Vespula xanthoptera*, is a potent antimicrobial peptide that interacts strongly with anionic liposomes [36]. The dissociation constant, *K*_D_, measured for a 1:1 mixture of neutral and anionic lipids is *K*_D_ = 15 nM, which ensures that all peptide is bound under the experimental conditions chosen here [36]. We measured the kinetics of content release by encapsulating the fluorescent dye CF at a high, self-quenching concentration. The interaction of MasX with the liposomes leads to dye efflux and the relief of dye self-quenching. The time course or fluorescence increase is measured, and the mean time constant of efflux, *τ*, determined. The mean time constant of efflux is a robust and model-independent observable. If efflux is measured at the same lipid and peptide concentrations, changes in τ can then be related to structural effects caused by alterations in the acyl chain compositions [50].

We measured dye release at room temperature and at 37 ^°^C, the growth temperature of the cultures from which the samples were isolated (Fig. 4).

**Figure 4.**
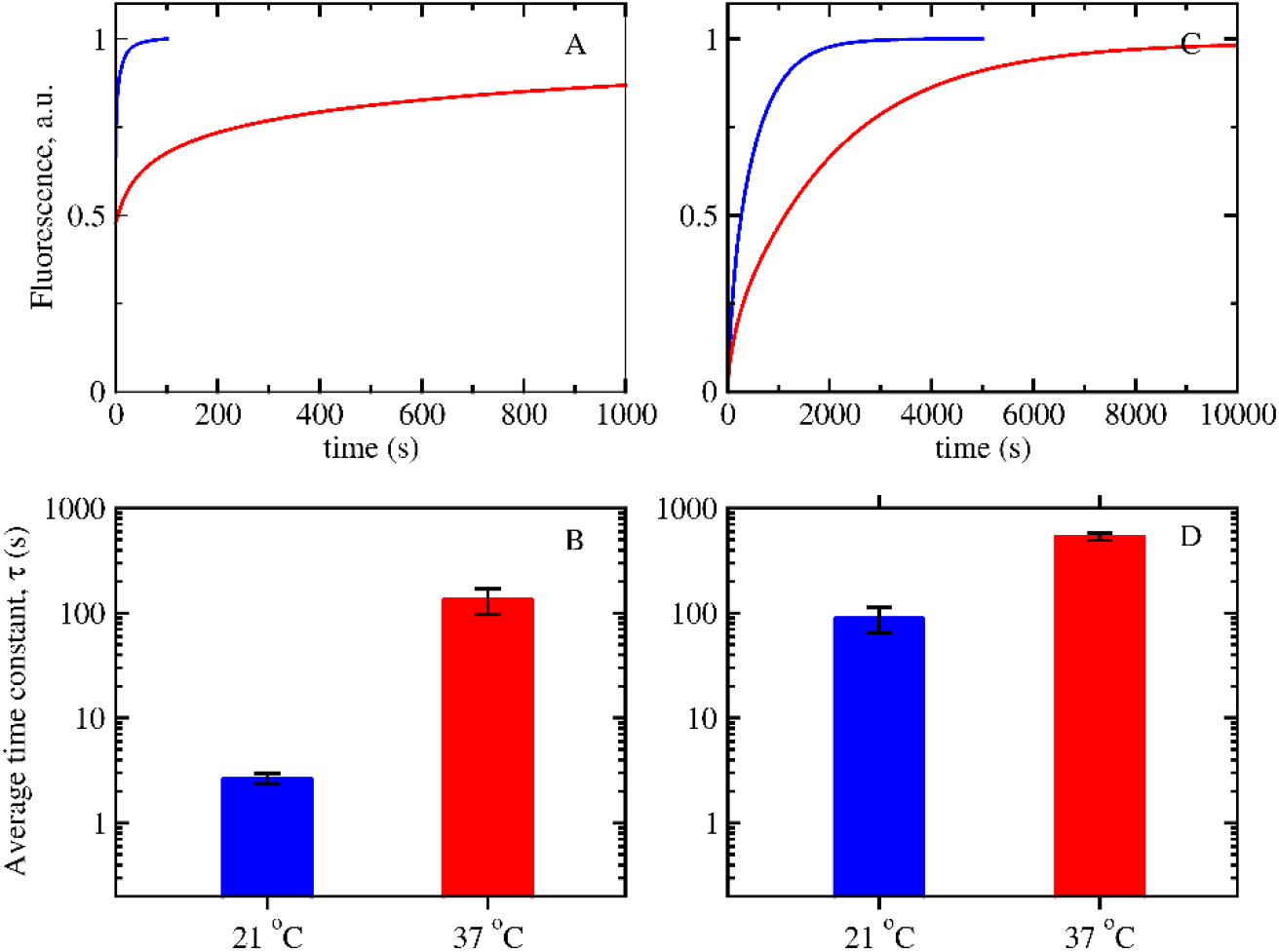
Kinetics of CF release induced by the antimicrobial peptide mastoparan X from large unilamellar lipid vesicles made from total lipid extracts. Aqueous peptide concentration was 5 μM in all experiments. Lipid phosphate concentration was 80 ± 20 μM. Representative dye release kinetics from liposomes based on lipid extracts from unsupplemented cultures (A), and from cultures supplemented with oleic acid (C). Room temperature, *blue trace;* 37 ^°^C, *red trace*. Average τ of dye release using liposomes based on lipid extracts from unsupplemented cultures (B), and from cultures supplemented with oleic acid (D) at two temperatures. Room temperature, *blue bars;* 37 ^°^C, *red bars*. Error bars represent the standard deviation from three independent samples.

#### 3.2.1 Kinetics of dye release: lipid extracts from cultures grown in TSB only

At 21 ^°^C, peptide-induced dye release from liposomes that were prepared using total lipid extracts from cultures grown in unsupplemented TSB occurred with a τ = 2.7 s (Fig. 4 A, B). This value is similar to that obtained from liposomes composed of 80:20 mixture of 1-palmitoyl-2-oleoylphosphatidylglycerol (POPG) and 1-palmitoyl-2-oleoylphosphatidylcholine (POPC). POPG and POPC (τ = 2.5 s, data not shown), under the same conditions. However, when the experimental temperature was raised to 37 ^°^C, peptide-induced dye efflux slows by a factor of 50 relative to that measured at room temperature (τ = 130 s). In contrast, for 80:20 mixture of POPG and POPC, the kinetics of dye release do not change significantly (τ = 3.4 s, data not shown).

#### 3.2.2 Kinetics of dye release: lipid extracts from cultures grown in TSB with oleic acid

Remarkably, the dye release measured using liposomes composed of total lipid extracts from oleic acid-supplemented cultures occurs very slowly. At 21 ^°^C, τ = 89 s (Fig. 4 C, D), and at 37 ^°^C, the kinetics slow down even more to τ = 510 s, which is significantly slower than that observed in TSB only liposomes, and two orders of magnitude slower than that measured in 80:20 mixture of POPG and POPC.

## 4. Discussion

It is well established that *S. aureus* and other bacteria adjust the lipid composition of their cytoplasmic membranes, depending on growth conditions [48,52–56]. Here, we sought to address more specifically the physico-chemical consequences of alterations in membrane lipid composition in response to growth medium supplementation by exogenous lipids. The focus of this study is on the phospholipid and glycolipid fractions, the two dominant lipid classes found in *S. aureus*.

The changes in membrane properties that occur because of the altered acyl chain profile in the glycolipid fraction from oleic acid-supplemented cultures are significant. When grown in standard TSB, the acyl chain profile of the phospholipid and glycolipid fractions are very similar [48]. In DSC scans of individual glycolipid and phospholipid fractions, the glycolipid fraction undergoes a melting transition shifted by about 15^°^C relative to the phospholipid fraction (Fig. 1C), as we would expect based on the strong headgroup interactions in glycolipids that stabilize the gel phase.

The addition of exogenous oleic acid to the medium leads to the appearance of odd-numbered, unsaturated species in both the glyco- and phospholipid fractions [48], indicating the incorporation of oleic acid into membrane lipids. For the phospholipid fraction, the distribution of species with the same total number of carbons is essentially the same with or without oleic acid supplementation [48]. However, in cultures grown in oleic acid-supplemented TSB, the acyl chain composition of the glycolipid fraction overall is shifted towards longer acyl chains relative to that from unsupplemented cultures (Fig. 5). The main DGDG species from non-supplemented cultures are 32:0 and 33:0. In the oleic acid-supplemented cultures, the distribution is shifted, with 35:1 and 35:0 becoming the major fractions. This increase in the total number of carbons from 33 to 35 appears subtle, but the consequences for the phase behavior of aqueous suspensions of glycolipid and whole lipid extracts are significant.

**Figure 5.**
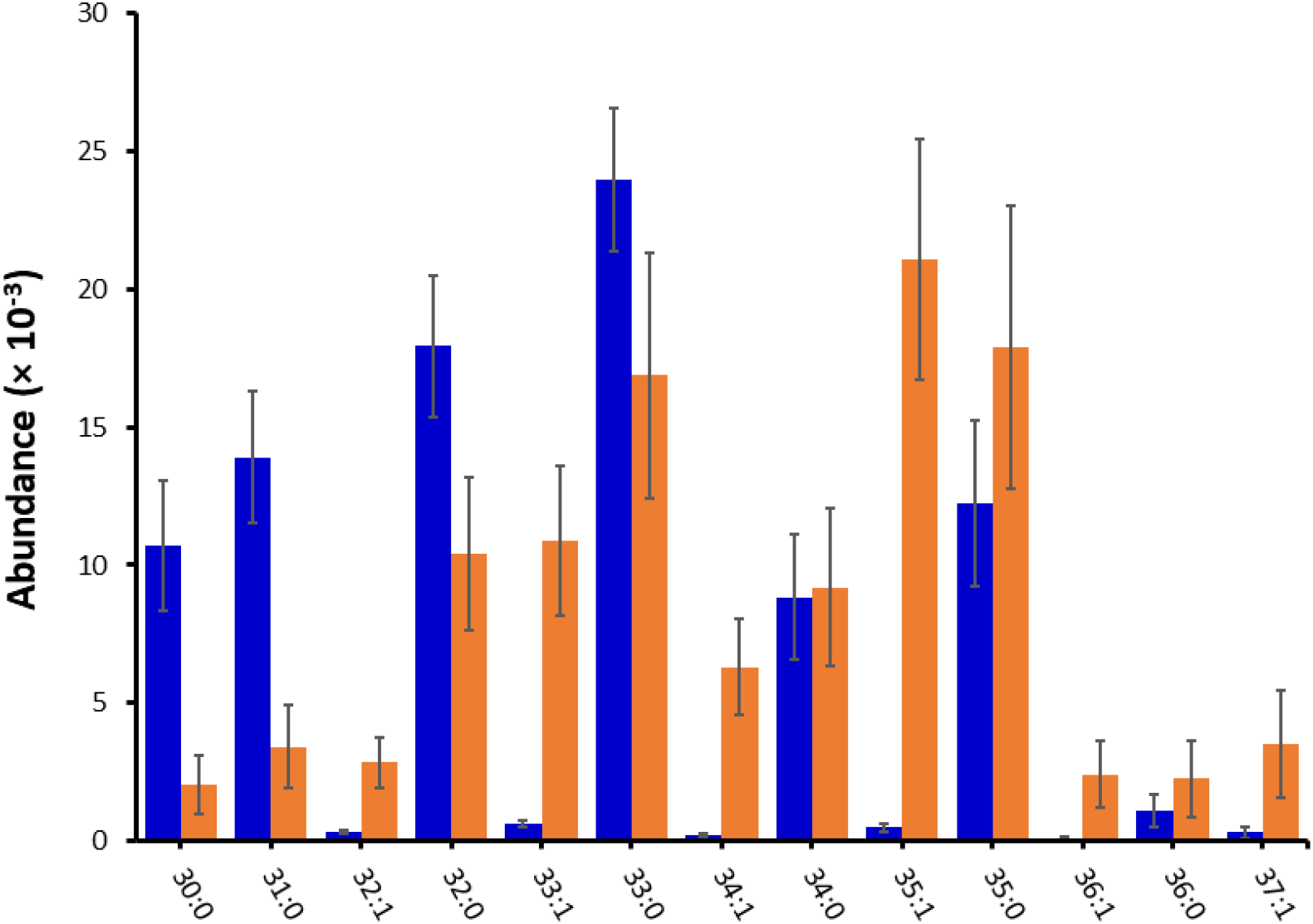
**Diglucosyldiglyceride (DGDG) species distribution** in extracts obtained from cultures grown in TSB, *blue bars* and from cultures grown in TSB supplemented with oleic acid, *orange bars*. Data was obtained from Hines et al. (2020) [48].

The polar headgroups in MGDG and DGDG are small compared to those of PG or phosphatidylcholine. The small headgroup size and the ability to establish hydrogen bonds between headgroups make glycolipids prone to adopt inverted hexagonal or cubic phases, rather than the flat, lamellar phases that are characteristic for biological membranes [23,57–59]. In inverted lipid phases, the polar headgroups are crowded in the interior of the assembly, enclosing an aqueous medium. The acyl chains are removed from contact with water, either by the formation of rods (inverted hexagonal phase) or a reticulated cubic phase [23,59] (Figure 6). The tendency to form non-bilayer phases for lipids with small polar headgroups depends strongly on the acyl chain composition, with longer and more unsaturated acyl chains promoting the formation of inverted phases with negative curvature [23,59].

**Figure 6.**
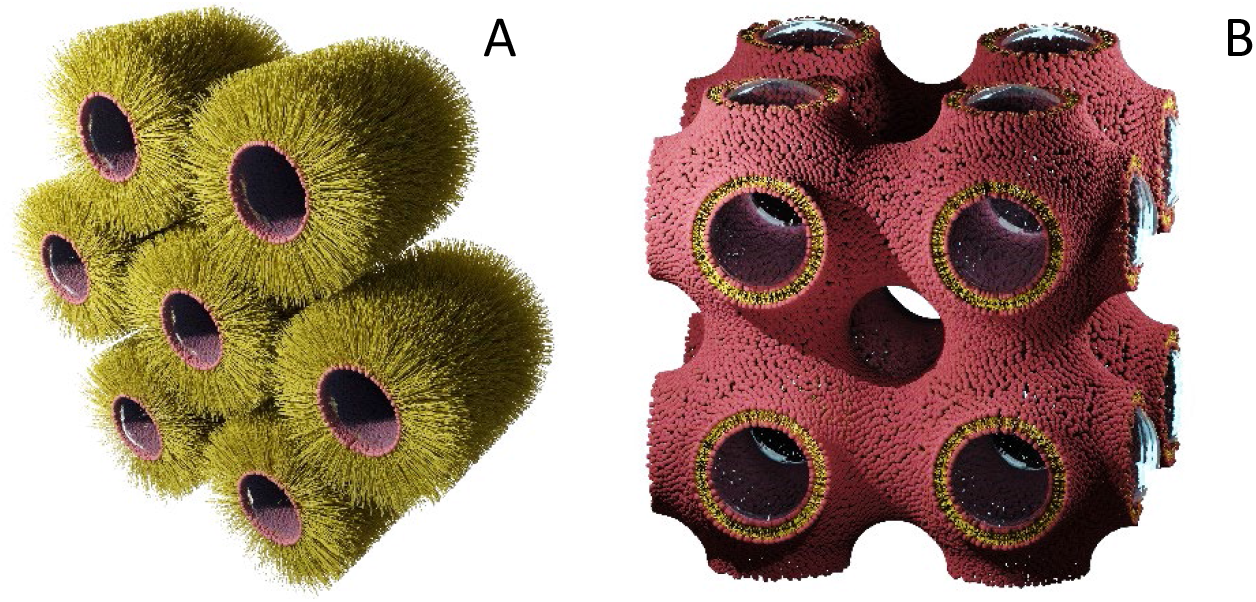
Artist’s representation of inverted lipid phases. Inverted hexagonal H_II_ (A) and an example of an inverted bicontinuous cubic phase (I*m*3*m*) (B). The drawings were generated using the *Blender* software [60].

We propose that the high temperature transition observed in the DSC scans on membrane preparations from cultures grown in oleic acid-supplemented medium is caused by the transition of a fluid lamellar phase to an inverted hexagonal phase, which is promoted by a higher proportion of C35 glycolipid species (35:0 and 35:1). It is important to note that, at the growth temperature, all membrane preparations investigated (fragments, whole lipid extracts, phospho- and glyco-lipid fractions) form a fully functional, fluid and flat lipid bilayer. We do not suggest the existence of inverted lipid phases at the growth temperature. Rather, the ability of the lipid extracts to transition from a lamellar to an inverted phase at temperatures above 60 ^°^C suggests that some degree of acyl chain packing frustration exists at the growth temperature. The packing frustration results from forcing lipids prone to form an inverted phase of negative spontaneous curvature into a flat lipid bilayer [44,45,50,61]. Whether the function of membrane proteins is altered as a consequence is beyond the scope of this work, but defect formation by the antimicrobial peptide MasX is clearly inhibited (Fig. 4).

Linear, α-helical antimicrobial peptides act by perturbing the cell membrane and forming transient or, more rarely, permanent defects. Their efficacy is in most cases inhibited by the presence of lipids that form inverted phases because peptide-induced membrane perturbation usually involves the formation of lipid structures with the opposite - positive – spontaneous curvature [50,62–68]. Thus, we attribute the slow dye efflux from liposomes constructed from whole lipid extracts from OA supplemented cultures (Fig. 4) to the presence of longer-chained glycolipid species that effectively protect the *S. aureus* cytoplasmic membrane against peptide-induced lysis.

Inverted hexagonal and cubic phases formed by synthetic lipids in excess water are well studied [23,57,59,69]. Symmetric monoglucosyl dialkyl and diacyl glycolipids with 32 total carbons or longer show a gel to inverted hexagonal phase transition at high temperatures [23]. Data on diglucosyl glycolipids are scarcer and limited to straight-chain, saturated species, which form lamellar phases even at high temperatures.

Inverted lipid phases have been observed in aqueous suspensions of lipid extracts from *Pseudomonas fluorescens* and the archaebacterium *Sulfolobus solfataricus* [70,71]. However, these are either rich in phosphatidylethanolamine (*Pseudomonas*) which is not present in *S. aureus*, or di- and tetra-alkyl ether lipids, unusual lipids characteristic for thermophile archaebacteria. The most comprehensive studies of inverted lipid phases in lipid mixtures of biological origin stem from lipid isolates of *Acholeplasma laidawii* [72–76]. *A. laidawii* responds to oleic acid supplementation with an increase in the fraction of DGDG relative to MGDG in the cytoplasmic membrane. This adaptation was thought to maintain an optimum balance between lipids with no spontaneous curvature that adopt lamellar phases, and those with spontaneous negative curvature. The result is a cytoplasmic membrane, in which the individual monolayers have a slightly negative spontaneous curvature to provide optimum stability [72–76].

In summary, oleic acid incorporation into the bacterial cytoplasmic membrane leads to a specific increase in long-chain saturated and unsaturated glycolipid species that are known to promote the formation of inverted lipid phases at temperatures well above the growth temperature. This observed shift in the acyl chain composition of the glycolipid fraction leads to an increased resistance to permeabilization by the antimicrobial peptide MasX at physiological temperature.

The uptake of exogenous oleic acid is clearly energetically advantageous for the organism but appears to simultaneously pose a challenge to the structural integrity of the cytoplasmic membrane. We do not know the exact nature of the structural consequences of oleic acid incorporation, but we may speculate that the enrichment in unsaturated lipid species leads to lipid packing defects and, consequently, a more permeable membrane. Alternatively, and perhaps more likely, the shift in the membrane lateral pressure profile upon incorporation of oleic acid may impair the function of integral membrane proteins that is restored to optimal functionality by the addition of long-chain glycolipids.

## 5. Acknowledgements

This work was supported by NIH grants 1R21AI13535 to BJW and CG, and RO1AI173144 to KMH and BJW.

